# An Integrative Boosting Approach for Predicting Survival Time With Multiple Genomics Platforms

**DOI:** 10.1101/338145

**Authors:** K. Y. Wong, C. Fan, M. Tanioka, J. S. Parker, A. B. Nobel, D. Zeng, D. Y. Lin, C. M. Perou

## Abstract

Recent technological advances have made it possible to collect multiple types of genomics data on the same set of patients. It is of great interest to integrate multiple genomics data types together for predicting disease outcomes. We propose a variable selection method, termed Integrative Boosting (I-Boost), that makes proper use of all available clinical and genomics data in predicting individual patient survival time. Through simulation studies and applications to data sets from The Cancer Genome Atlas, we demonstrate that I-Boost provides substantially higher prediction accuracy than existing variable selection methods. Using I-Boost, we show that (1) the integration of multiple genomics platforms with clinical variables significantly improves the prediction accuracy for survival time over the use of clinical variables alone; (2) gene expression values are typically more prognostic of survival time than other genomics data types; and (3) gene modules/signatures are at least as prognostic as the collection of individual gene expression data.

## Background

Prediction of disease outcomes, such as individual patient survival time, is critically important for cancer patients. Traditional prognostic models that rely solely on clinical variables, such as age and tumor stage, fail to account for the molecular heterogeneity of tumors and thus may lead to suboptimal treatment decisions [1]. To remedy this situation, many studies have incorporated gene expression data in survival prediction [2, 3, 4, 5].

Large-scale genomics projects such as The Cancer Genome Atlas (TCGA) have generated detailed molecular data on patients with a variety of cancer types. In TCGA, six types of “omics” data have been collected on the same set of patients: DNA copy number variation, somatic mutation, mRNA expression, microRNA expression, DNA methylation, and expression of ~200 proteins/phosphoproteins. The availability of multiple data types has enabled researchers to address a variety of important questions. For example, patients can be more precisely classified into molecular subtypes based on integrative clustering of multiple genomics data types or platforms [6, 7, 8]. In addition, it is possible to identify genes that are related to patient survival time by decomposing the expression of each gene into a component that is explained by the methylation level and a component that is not [9].

One unsolved issue in cancer genomics is the prognostic value of integrated genomics and clinical data versus clinical data only. Yuan et al. [10] compared models with clinical data only versus models with both clinical and genomics data on various cancer types and concluded that genomics data provide only a limited gain in survival prediction accuracy. In their analysis, however, potential differences among data types were not taken into account. For breast cancer, for instance, the combination of genomics and clinical data has been shown to improve outcome predictions [11, 12]. A major goal of the present work is to fully explore the predictive power of integrating clinical and genomics data together.

A second unsolved issue is the prognostic value of individual gene expression values (~25,000) versus a predefined set of gene expression signatures or “modules” (~500). Gene modules have been developed for representing distinct cell types (e.g., epithelial, immune, endothelial, etc.), specific biological processes, or activated molecular signaling pathways. They have been shown to successfully capture signaling pathway activitives or cell type heterogeneity within tumors. We wish to investigate whether individual gene expression data or existing gene modules provide more accurate outcome prediction.

A third unsolved issue is the relative importance of different types of genomics data in outcome prediction. Different data types are collected at different costs and also with widely varying feature spaces. Naturally, not all data types are equally important in outcome prediction. We aim to determine which data types may be omitted from analysis without a significant reduction in prediction accuracy.

An overarching methodological challenge in addressing the aforementioned issues is the identification of genomic variables predictive of survival time when the number of variables is much larger than the sample size. Penalized regression methods, such as least absolute shrinkage and selection operator (LASSO) [13] and elastic net [14], are commonly used to identify important genomic variables. When variables are highly correlated, elastic net tends to have better performance in prediction than LASSO [14]. However, both LASSO and elastic net are generic variable selection procedures that do not distinguish different types of data and thus tend to select more variables from the data types with larger numbers of variables. Because different data types capture different biological structures, both large and small data types may carry important signals. Methods that treat all variables equally may not be able to pick out independent signals from small data types. In addition, LASSO and elastic net impose the same penalty on all regression parameters, which may be overly restrictive because the number of variables and the signal strength vary drastically across data types.

Boosting is an alternative to penalization for model estimation and prediction in high-dimensional settings. It was originally developed for binary classification in machine learning [15, 16]. The idea of boosting is to iteratively reweight the observations, with larger weights given to observations that are misclassified at the previous iteration, and apply simple classifiers on the reweighted data; their results are then combined to produce an aggregated classification procedure. Boosting was later generalized as a forward stagewise additive modeling method for statistical estimation [17, 18], which can be applied to many problems, including regression analysis for survival data [19]. Because of its flexibility in modeling choices and stability in high-dimensional settings, boosting has found applications in genomics studies; see the references in Mayr et al. [20, 21]. As in the case of LASSO and elastic net, however, existing boosting methods, such as component-wise boosting [22], do not distinguish variables of different data types.

To overcome the limitations of LASSO, elastic net, and existing boosting methods, we develop a novel method, termed Integrative Boosting (I-Boost), which combines elastic net with boosting. In I-Boost, the prediction rule is constructed iteratively, where at each iteration, the predictive power of each data type (conditional on the current prediction rule) is evaluated separately and the most predictive data type is selected to update the prediction rule using elastic net. Thus, independent signal from each data type can be incorporated into the prediction rule, and small but predictive data types will not be dominated by data types with large numbers of variables. In addition, the penalties on the regression parameters are learned data-adaptively and separately for different data types. Herein, we demonstrate the advantages of I-Boost using simulation studies and empirical data from the TCGA on patients with eight different cancer types. More importantly, we use I-Boost to address the aforementioned three unsolved issues in cancer genomics.

## Results and discussion

### Background

Suppose that there are *K* types of clinical or genomics predictors, with *d_k_* components for the *k*th type (*k* = 1,…, *K*). For *k* = 1,…, *K*, let ***X*** denote the *d_k_*-vector of predictors of the *k*th type. Write ***X*** = (***X***^(1)T^,…, ***X***^(*K*)T^)^T^. Let *T* denote the survival time of interest. We relate *T* to ***X*** through the proportional hazards model [23], such that the conditional hazard function of *T* given ***X*** takes the form of *h*_0_(*t*)exp(*β*^T^***X***), where *h*_0_(*t*) is an arbitrary baseline hazard function, *β* = (*β*^(1)T^,…, *β*^(*K*)T^)^T^, and *β*^(*k*)^ is a *d_k_*-vector of regression parameters associated with ***X***^(*k*)^.

The survival time *T* is subject to right censoring by *C*, such that we observe *Y* ≡ min(*T, C*) and Δ ≡ *I*(*T* ≤ *C*), where *I*(·) is the indicator function. For a study with *n* patients, the data consist of (*Y_i_*, Δ_*i*_, ***X***_*i*_) (*i* = 1,…, *n*). The partial likelihood [24] for *β*
is

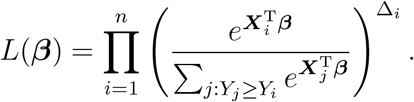

### LASSO and elastic net

Because ***X*** is high-dimensional, it is not feasible to estimate *β* by maximizing the partial likelihood. One possible remedy is to impose sparsity assumptions on *β* and adopt penalization methods, such as LASSO [13] and elastic net [14]. LASSO estimates *β* by maximizing the *L*_1_-penalized log-partial likelihood function

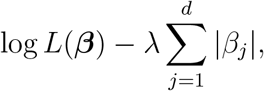

where 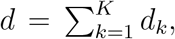, and λ is a tuning parameter. Elastic net generalizes LASSO by including an *L*_2_ penalty, such that the objective function becomes

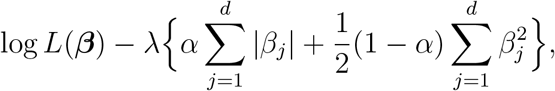

where *α* ∈ [0, 1] is a tuning parameter that controls the relative magnitudes of the *L*_1_ and *L*_2_ penalties. (When *α* =1, elastic net reduces to LASSO.) The implementation of LASSO and elastic net is described in Methods.

For both LASSO and elastic net, the penalty term dominates under large values of λ, and the parameter estimates tend to be small with some values being exactly zero. Unlike LASSO, elastic net exhibits the grouping effect in that the regression parameters for a group of highly correlated variables tend to be equal, which is desirable in the context of gene selection [14]. Both LASSO and elastic net impose the same penalization on each regression parameter and thus do not distinguish different types of predictors. As a result, these methods may be inefficient when certain data types are much more predictive than others.

### I-Boost

To account for the differential predictive power of different data types, we propose a boosting algorithm called I-Boost. Boosting is an iterative optimization algorithm that minimizes a loss function 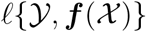 over a class of functions of predictors 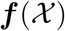, where 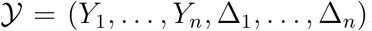, 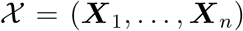, and 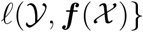 measures the deviation of the prediction 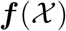 from the outcome 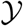. At each iteration, we update 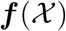 additively by the value 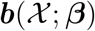 up to a scaling factor, where ***b*** is a fixed basis function, and *β* is a vector of parameters. Specifically, at the *m*th iteration, we find *β*^(*m*)^ that minimizes 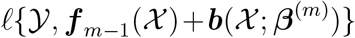, possibly under some constraints on *β*^(*m*)^, where ***f***_*m*–1_ is the estimate of ***f*** at the (*m* – 1)th iteration. Then, we set 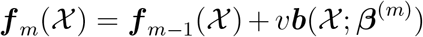 for some fixed step-length factor *v* ∈ (0, 1]. We terminate the iterations when some stopping criterion is satisfied.

In I-Boost, we set the loss function 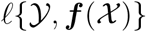 to be the negative log-partial likelihood function and the basis function to be 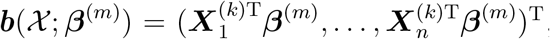 where 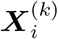 is the vector of the *k*th type of predictors for the *i*th patient, and the data type *k* is selected separately at each iteration based on the values of the loss function obtained using different data types. To handle high-dimensional data, we impose an elastic-net penalty on *β*^(*m*)^ in the optimization step. Effectively, we perform maximum penalized log-partial likelihood estimation with an offset term 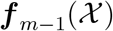 using a single data type at each iteration. Unlike existing boosting methods, such as component-wise boosting, the basis function in our case is a function of all variables of a data type instead of a single variable. This choice of basis function is motivated by the expectations that some data types are much more predictive than others and that the inclusion of less predictive data types may reduce the prediction accuracy of the model. By considering each data type separately, we perform selection on the data-type level at each iteration.

We propose two versions of I-Boost, namely I-Boost-CV and I-Boost-Permutation, which use cross validation and permutation, respectively, to choose the tuning parameters of elastic net at each iteration. The permutation procedure randomly permutes the outcome variables in order to remove association between the predictors and the outcome, and the tuning parameters are chosen such that no predictor is selected in half of the permuted data sets. The procedures are described in detail in Methods.

### Simulation studies

We conducted simulation studies to evaluate the performance of LASSO, elastic net, and the two versions of I-Boost. We considered three simulation settings, with different distributions of signals across the data types. In all three settings, a relatively large proportion of the signals is contributed by the clinical variables. The distributions of signals are shown in Figure 1, and the details of the simulation settings are provided in Methods.

**Figure 1.**
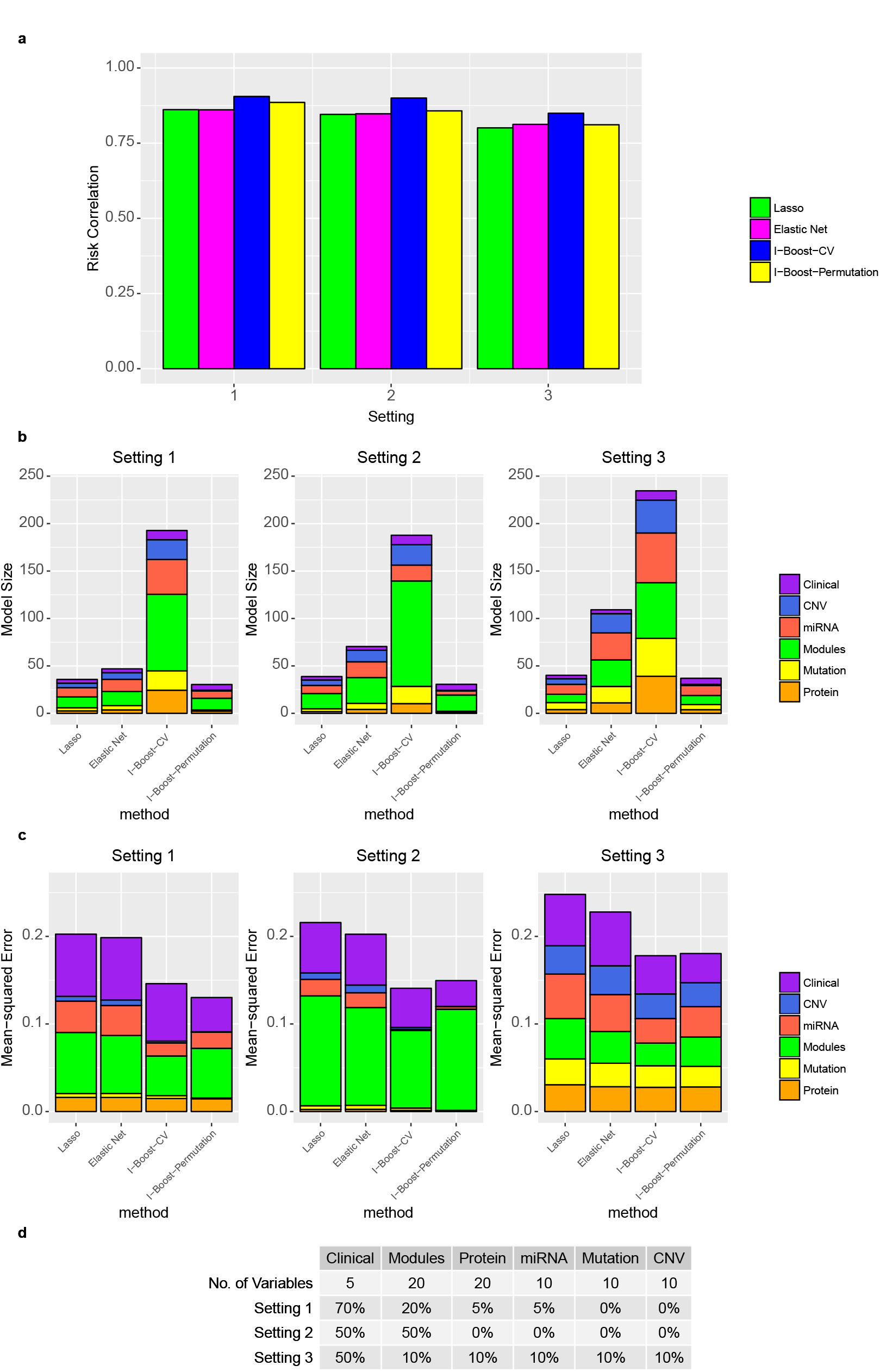
Simulation settings and results. **a** Prediction accuracy of LASSO, elastic net, I-Boost-CV, and I-Boost-Permutation measured by risk correlation under three different settings. **b** The average number of variables selected by the four methods under three different settings. Different types of the selected variables are represented by different colors. **c** Mean-squared error of the four methods under three different settings. The error is decomposed into errors of parameters for different data types, as represented by different colors. **d** Number of signal variables and distribution of signals across different data types for the three simulation settings. The number of signal variables is zero if the proportion of signals of the data type is 0%. Abbreviations are as follows: GeneExp represents individual gene expression; Module represents gene module; Clinical represents clinical variable; CNV represents copy number variant; Mutation represents somatic mutation; miRNA represents micro-RNA expression; and Protein represents protein expression.

We assessed the performance of the methods by the quality of prediction and parameter estimation. For prediction, we report the correlation between the estimated risk score 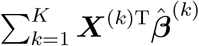 and the true risk score 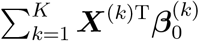, where 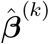 and 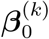 are the estimated and true parameter vectors, respectively. A higher correlation represents a greater degree of agreement between the predicted and actual outcomes. We call this measure the risk correlation. For parameter estimation, we report the mean-squared error 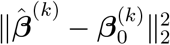 for *k* = 1,…, *K*.

Figure 1 shows the risk correlation and mean-squared error for elastic net, LASSO, and the two versions of I-Boost based on 1,000 replications; the average number of variables selected for each data type is also shown. The results show that I-Boost-CV always selects the largest number of variables, followed by elastic net, LASSO, and I-Boost-Permutation. In particular, I-Boost-CV tends to select a relatively large number of variables from data types with very weak or no signal. Nevertheless, evaluation of the mean-squared error reflects that the magnitude of the estimated regression parameters for those data types is small. In fact, both versions of I-Boost yield more accurate estimation than LASSO and elastic net in all cases.

For prediction, the two I-Boost methods perform the best overall. In all settings, I-Boost-CV produces more accurate prediction than all other methods. In Settings 1 and 2, where most signals are concentrated on only one or two data types, I-Boost-Permutation produces more accurate prediction than both elastic net and LASSO. In Setting 3, I-Boost-Permutation performs similarly to elastic net, while LASSO performs worse than I-Boost-Permutation. Between the two versions of I-Boost, I-Boost-CV tends to yield better prediction than I-Boost-Permutation, possibly because of the larger number of variables selected by I-Boost-CV. Thus, if the main interest is the selection of relevant variables, then one might consider I-Boost-Permutation for more conservative variable selection, even though this method is somewhat inferior in prediction when compared to I-Boost-CV.

We implemented LASSO, elastic net, and the two versions of I-Boost using R-3.2.2 on a 2.93 GHz Xeon Linux computer. On average, performing LASSO, elastic net, I-Boost-Permutation, and I-Boost-CV on one simulated data set (that consists of 500 subjects, 6 data types, and 1,294 predictors) takes about 2 minutes, 14 minutes, 3 hours, and 38 hours, respectively. I-Boost-CV is computationally intensive because in each iteration, cross validation is conducted on a three-dimensional grid. By contrast, in each I-Boost-Permutation iteration, the tuning parameter α is fixed at 1, no cross validation is involved in the selection of λ, and LASSO is performed only once for each data type. Therefore, I-Boost-Permutation may serve as a computationally efficient alternative to I-Boost-CV.

### Evaluation of LASSO, elastic net, and I-Boost using TCGA data

We next evaluated the performance of the methods using three TCGA data sets, namely the lung adenocarcinoma (LUAD) data set, the kidney renal clear cell cancer (KIRC) data set, and a pan-cancer data set derived from ~1,400 patients that represents eight different tumor types considered by Hoadley et al. [25]; see Methods for a detailed description of the data sets and the evaluation procedure. For each data set, we first split the data 30 times into training and testing sets. We then performed LASSO, elastic net, and the two versions of I-Boost for various combinations of data types on patients from the training set of each split. For each combination of data types and each split, we calculated the risk scores for patients in the testing set using the estimates from the corresponding training set, and we used the concordance index (C-index) [26] to evaluate the prediction accuracy of the risk scores.

The average C-index values over the splits obtained from LASSO and elastic net are given in Figure 2. For the KIRC and pan-cancer data sets, the prediction tends to be much better than random (i.e., the C-index values are much larger than 0.5). For the LUAD data set, which has a small sample size, some of the models yield relatively poor prediction (with C-index values smaller than 0.6). For many models, the predictive performance of elastic net is either similar or superior to LASSO.

**Figure 2.**
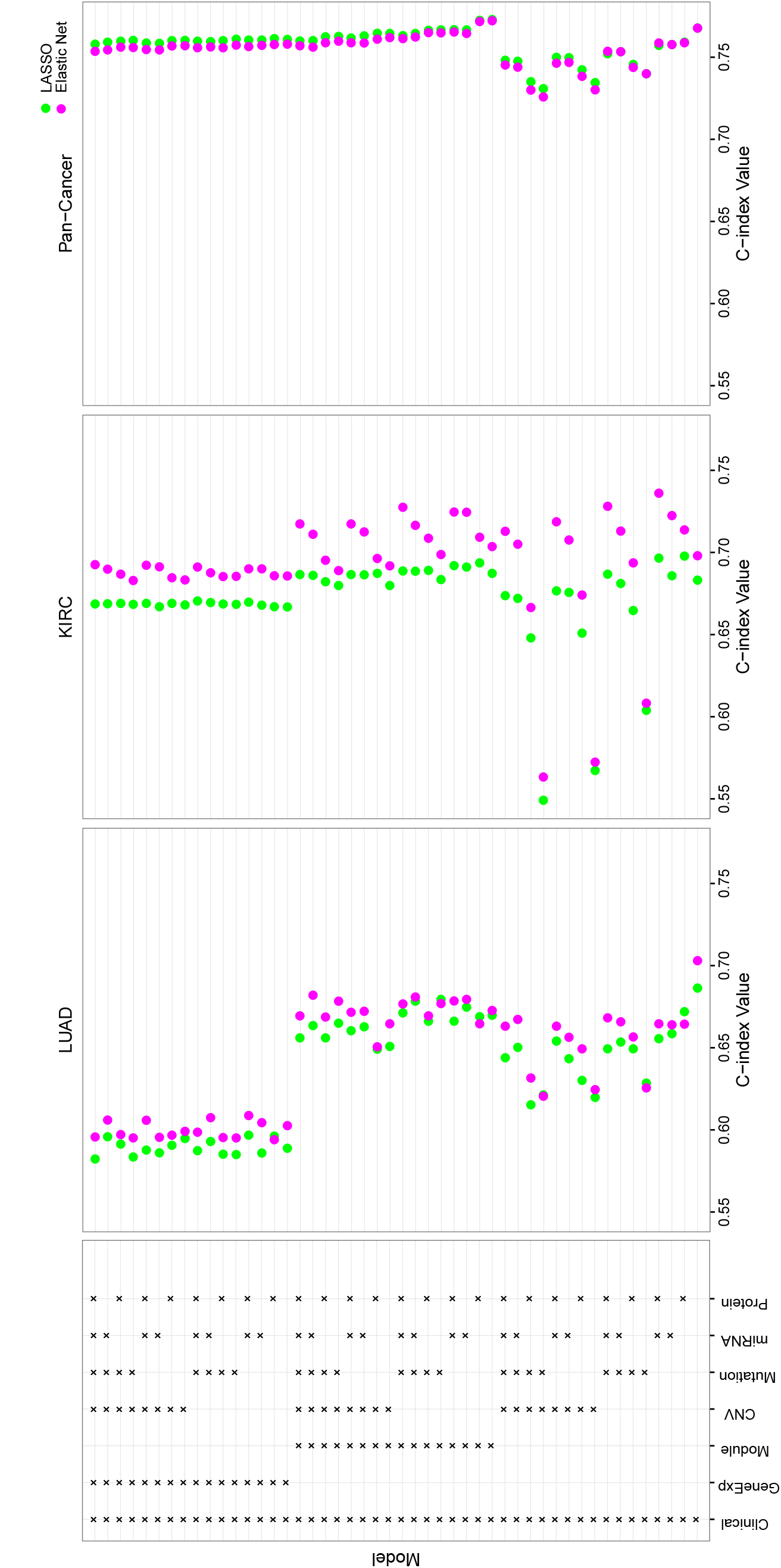
Analysis results for the TCGA LUAD, KIRC, and pan-cancer data sets using LASSO and elastic net. Each row represents a particular combination of data types used as predictors, as indicated by the box on the left. Each dot is an average C-index value obtained by performing LASSO or elastic net on 30 training and testing data set pairs. See the caption of Figure 1 for the abbreviations of the data types.

For LASSO and elastic net, the models containing more data types as predictors do not necessarily perform better than those with fewer data types. One possible explanation is that the extra data types may contain very little relevant information on patient survival, such that adding those data types introduces more noise than signal into the model. In practice, however, it is challenging to decide which data types to consider without prior knowledge of their importance.

Figure 3 shows the average values of the C-index obtained from elastic net, I-Boost-CV, and I-Boost-Permutation for different models. For the LUAD, KIRC, and pancancer data sets, both versions of I-Boost provide better prediction than elastic net in almost all cases. The difference in prediction accuracy between I-Boost and elastic net is particularly large when the sample size is small and the number of predictors is large. The difference is likely due to the fact that I-Boost involves the selection of data types, so that the large and non-predictive data types would not be selected in most iterations, and their presence would not substantially worsen the prediction accuracy. For the KIRC and pan-cancer data sets, I-Boost-CV yields better prediction than I-Boost-Permutation, whereas for LUAD, there are no clear differences between the two methods.

**Figure 3.**
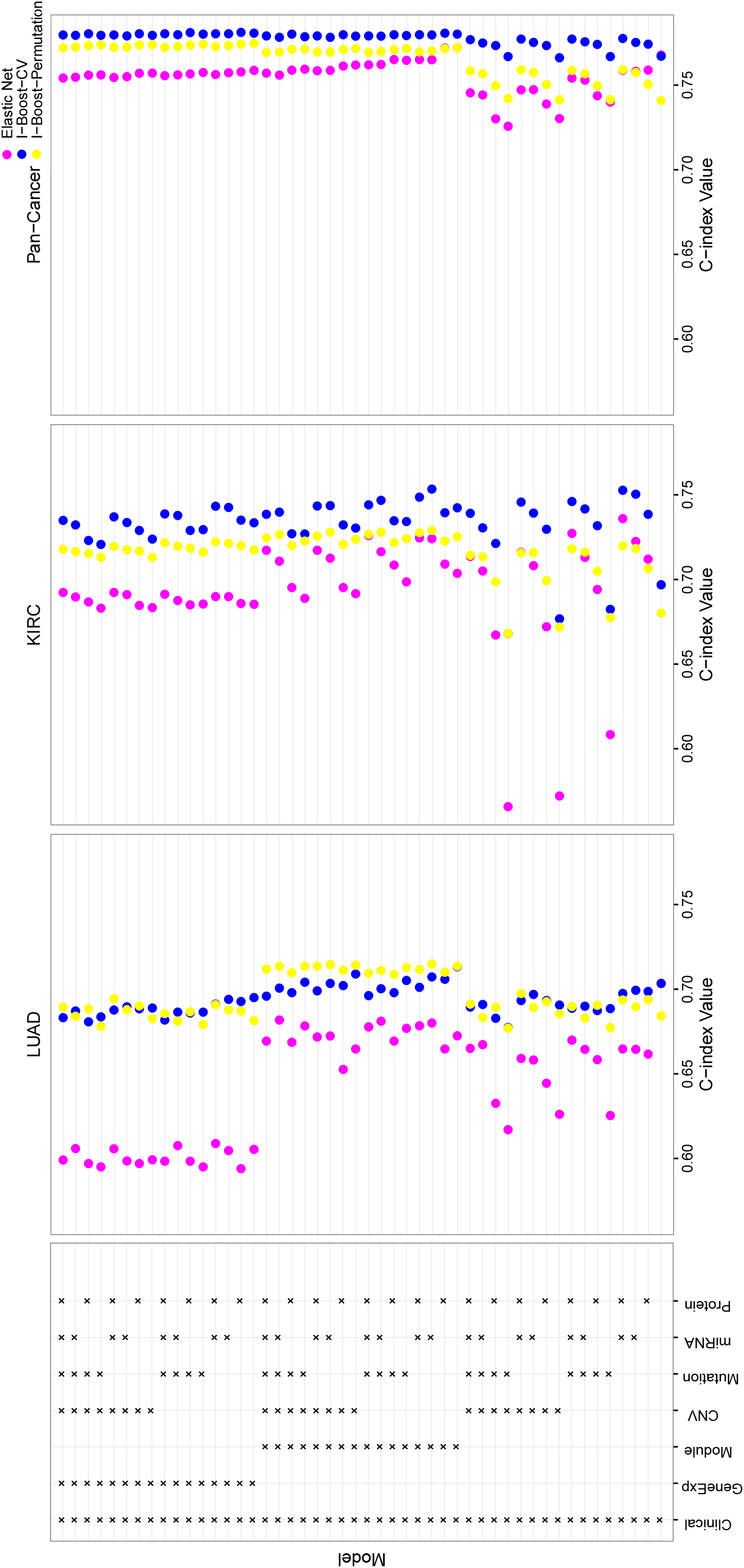
Analysis results for the TCGA LUAD, KIRC, and pan-cancer data sets using elastic net, I-Boost-CV, and I-Boost-Permutation. Each row represents a particular combination of data types used as predictors, as indicated by the box on the left. Each dot is an average C-index value obtained by performing elastic net, I-Boost-CV, or I-Boost-Permutation on 30 training and testing data set pairs. See the caption of Figure 1 for the abbreviations of the data types.

### Prognostic value of integrated clinical and genomics data

To assess whether the genomic variables provide extra predictive power in the presence of the clinical variables, we compared the values of the C-index obtained by I-Boost-CV or I-Boost-Permutation under the model with clinical variables only and the models with both clinical and genomic variables. The plots of the C-index values are provided in Figure 4. For reference, we also computed the C-index from the conventional maximum partial likelihood estimation for the model with clinical variables only.

**Figure 4.**
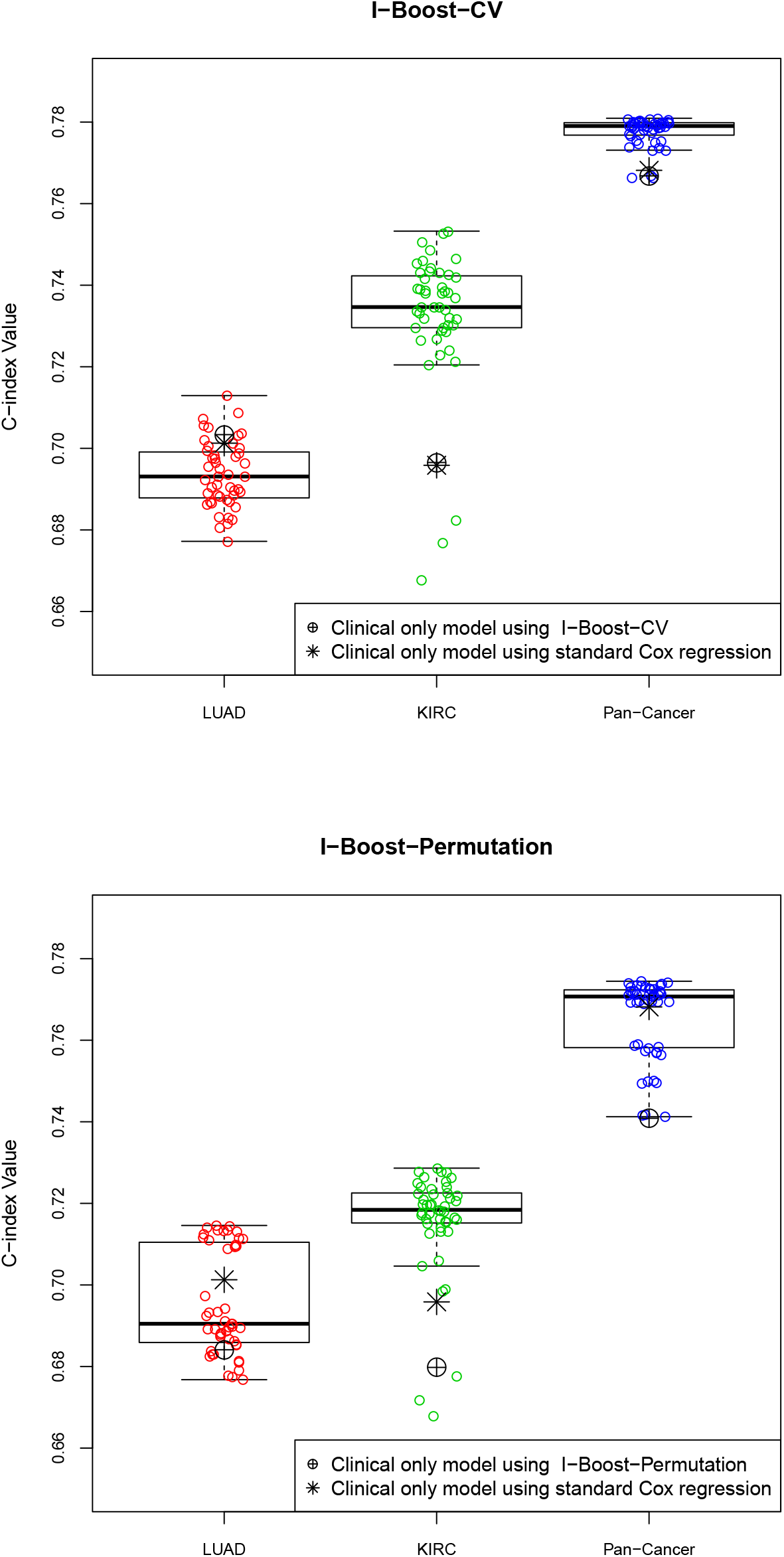
C-index values for the TCGA LUAD, KIRC, and pan-cancer data sets using I-Boost-CV or I-Boost-Permutation. Each dot represents the average C-index value obtained by performing I-Boost-CV or I-Boost-Permutation on a set of predictors (that contains the clinical variables) over 30 training and testing data set pairs. The average C-index values obtained by fitting I-Boost-CV, I-Boost-Permutation, or the standard Cox regression on the clinical variables are marked.

The patterns of the results from I-Boost-CV and I-Boost-Permutation are similar. For the KIRC and pan-cancer data sets, the majority of the models that contain both clinical and genomic variables provide better prediction than either I-Boost or maximum partial likelihood estimation with clinical variables only; the difference in prediction accuracy is particularly large under I-Boost-CV. For the LUAD data set, only a few models that contain both clinical and genomic variables provide better prediction than the model with clinical variables only for both I-Boost-CV and I-Boost-Permutation. These results indicate that in certain cancer types, genomic variables contribute to survival prediction in the presence of clinical variables, and the magnitude of the contribution can be large. When the same comparisons are made using LASSO or elastic net, however, the inclusion of genomic variables in the models does not appreciably improve prediction.

### Evaluation of gene expression modules

To compare the performance of gene modules versus individual gene expression data, we calculated the C-index values for models with each type of gene expression data separately. Specifically, for each combination of data types other than individual gene expression data and gene modules, we computed the difference between the C-index values obtained from I-Boost-CV or I-Boost-Permutation on those data types with gene modules and on those data types with individual gene expression data. The differences in the C-index are shown in Figure 5. Under both methods, the use of gene modules leads to better prediction than the use of expression data of all individual genes for the LUAD and KIRC data sets. For the pan-cancer data set, the performance of gene modules is slightly worse than that of individual gene expression data. Nevertheless, the differences in C-index values are small in all cases, and there is no strong evidence favoring gene modules or individual gene expression data on the basis of prediction accuracy. Because gene modules are smaller in number and much easier to interpret, we recommend the use of gene modules over individual gene expression data.

**Figure 5.**
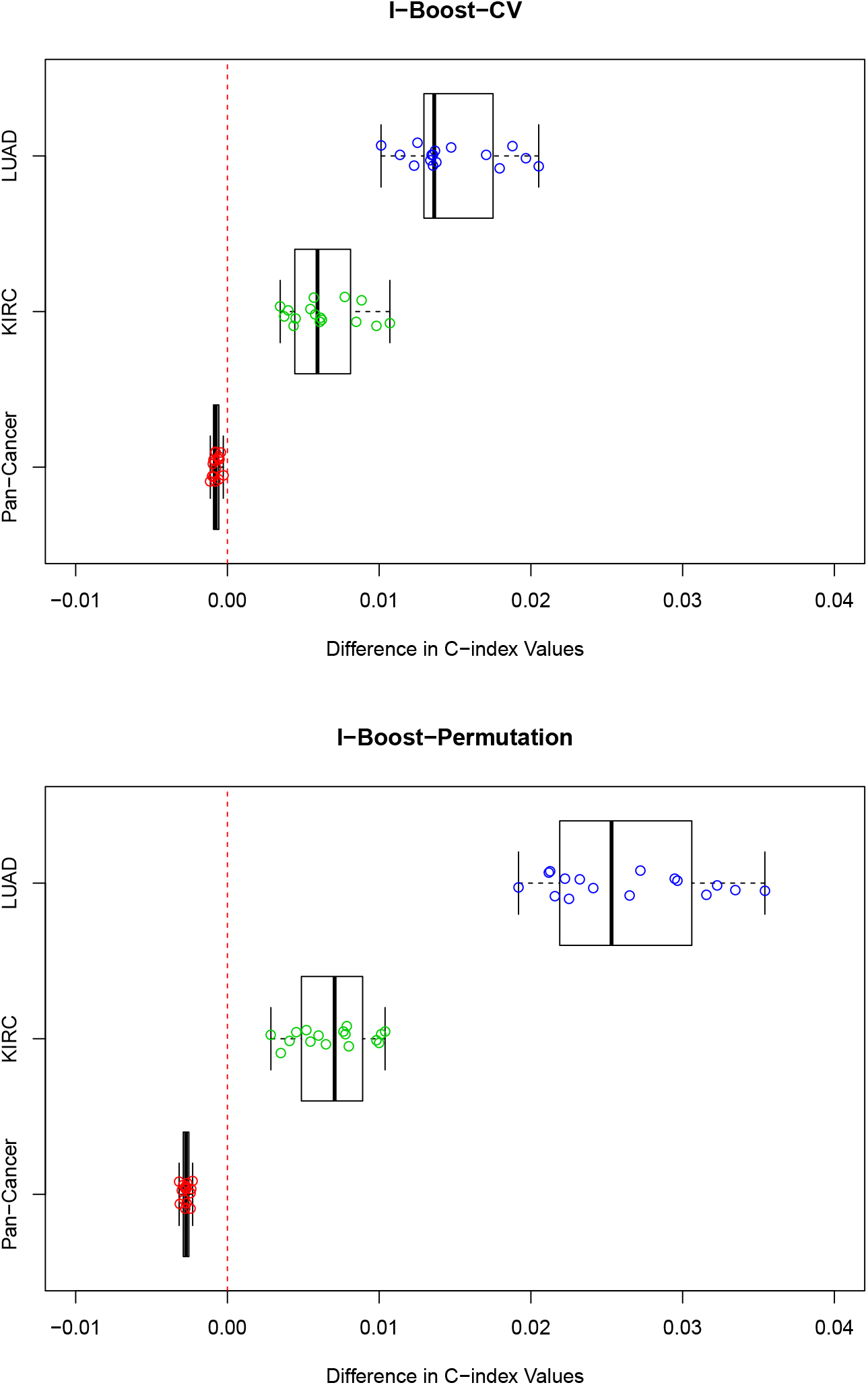
Comparison of C-index values for models containing individual gene expression data and models containing gene modules using the TCGA LUAD, KIRC, and pan-cancer data sets. Each dot represents the difference in average C-index values obtained by fitting I-Boost-CV or I-Boost-Permutation on two sets of predictors over 30 training and testing data set pairs. The first set of predictors contains a combination of data types and gene modules; the second set of predictors contains the same combination of data types and individual gene expression data. A positive difference represents better prediction using the model with gene modules.

### Comparison among genomics data types

To evaluate the relative prognostic value of each genomics data type, we formed a series of nested models as follows. We began with the model containing clinical variables only. Then, we compared the models containing clinical variables and a type of genomic variables and selected the most predictive model. This process was repeated until all data types were included, and the model selected at each step contained all of the previously selected data types. Individual gene expression data were not considered in this analysis. The order in which the genomics data types entered the models reflects their relative importance. We performed this procedure for elastic net and the two versions of I-Boost. For the LUAD, KIRC, and pan-cancer data sets, the C-index values for the series of models are plotted in Figure 6, and the data type selected at each step is shown. We also plotted the average number of variables selected for each model.

**Figure 6.**
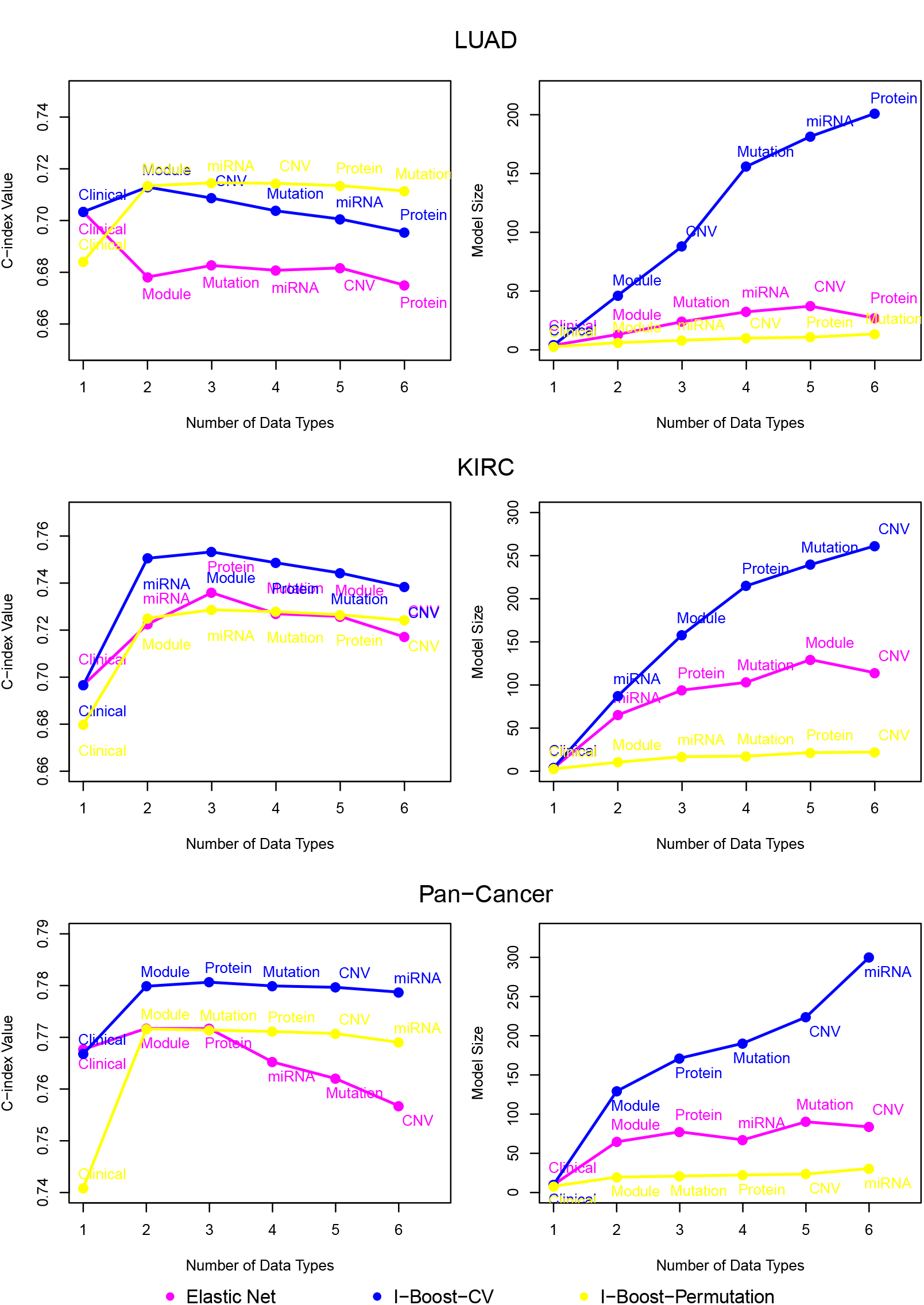
Analysis results for the TCGA LUAD, KIRC, and pan-cancer data sets, using elastic net, I-Boost-CV, and I-Boost-Permutation on nested models. In the left panel, each dot represents the average C-index value obtained by fitting elastic net, I-Boost-CV, or I-Boost-Permutation over 30 training and testing data set pairs. The leftmost dot represents the average C-index value for the model with clinical variables only. Each of the other dots represents the largest average C-index value among models that contain one more data type than the model corresponding to the dot on the left. In the right panel, the average number of selected variables for the models shown in the left panel is plotted. For both panels, beside each dot, the name of the additional data type is included. See the caption of Figure 1 for the abbreviations of the data types.

For the LUAD, KIRC, and pan-cancer data sets, the C-index under I-Boost-CV or I-Boost-Permutation tends to increase or stay approximately the same with the inclusion of each new data type. This indicates that I-Boost extracts useful information from each additional data type and that its performance tends not to be worsened by the inclusion of additional variables. By contrast, there is no clear improvement in prediction accuracy for elastic net when more data types are included.

I-Boost-Permutation always selects the smallest number of variables, followed by elastic net and I-Boost-CV. This finding is consistent with the conclusions from the simulation studies. Because the C-index obtained by I-Boost-Permutation is higher in most cases than that obtained by elastic net, we conclude that I-Boost-Permutation provides the same or better prediction using fewer variables than elastic net.

For the LUAD and pan-cancer data sets, gene modules are the first genomics data type selected under both versions of I-Boost, and the inclusion of gene modules leads to considerable improvement in prediction accuracy. For the KIRC data set, miRNA expression data are first selected by I-Boost-CV, while gene modules are first selected by I-Boost-Permutation. Nevertheless, for I-Boost-CV, the model with clinical variables and gene modules yields a C-index of 0.742, which differs only by 0.009 compared to the C-index of the model containing clinical variables and miRNA expression. For both versions of I-Boost, after the inclusion of the first genomics data type, the improvement in prediction accuracy with the inclusion of additional data types is marginal. We conclude that gene modules are overall the most predictive genomics data type, and the remaining genomics data types tend not to provide extra predictive power beyond clinical variables and gene modules.

### Important predictors for the LUAD, KIRC, and pan-cancer data sets

To obtain the final models of important predictors, we performed I-Boost-Permutation on the LUAD, KIRC, and pan-cancer data sets. The final models are shown in Tables 1–3 for the LUAD, KIRC, and pan-cancer data sets, respectively.

**Table 1:**
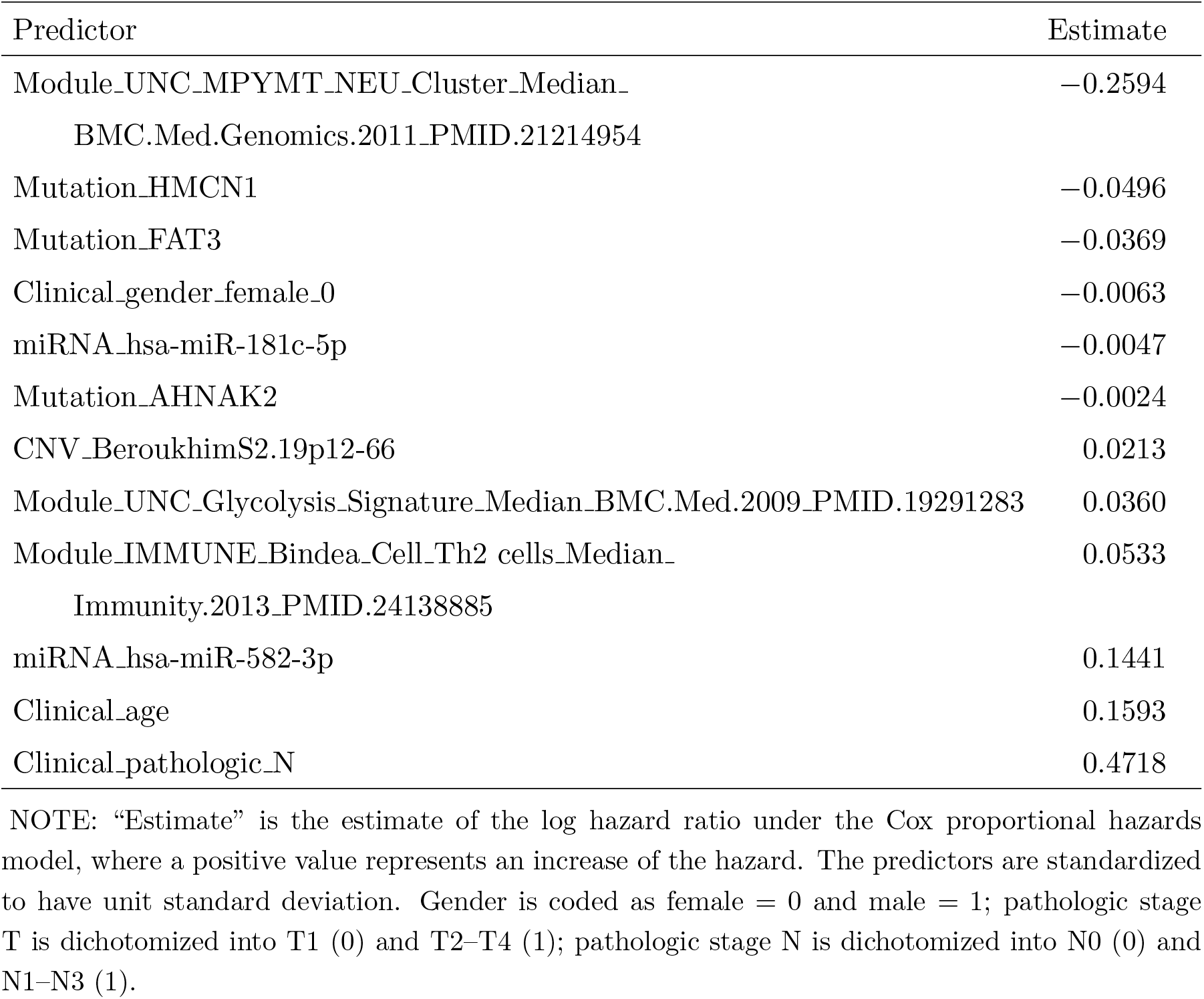
Analysis results from I-Boost-Permutation for the TCGA LUAD data set.

**Table 2:** Analysis results from I-Boost-Permutation for the TCGA KIRC data set.

| Predictor | Estimate |
| --- | --- |
| Protein_AR | -0.1076 |
| Module_IMMUNE_Bindea_Cell_CD8 T cells_Median_<br>Immunity.2013_PMID.24138885 | -0.0738 |
| Module_Mature_LuminaUp_Median_Nat.Med.2009_PMID.19648928 | -0.0669 |
| Module_UNC_MM_Red2_Median_BMC.Med.Genomics.2011_PMID.21214954 | -0.0648 |
| Module_GP7_Estrogen signaling: r=0.97 | -0.0571 |
| Module_UNC_HS_Green1_Median_BMC.Med.Genomics.2011_PMID.21214954 | -0.0396 |
| miRNA_hsa-miR-10b-3p | -0.0337 |
| miRNA_hsa-miR-192-5p | -0.0290 |
| Protein_Src_pY416 | -0.0288 |
| Module_UNC_LUMINAL_Cluster_Median_<br>BMC.Med.Genomics.2011_PMID.21214954 | -0.0162 |
| Module_UNC_HS_Green8_Median_BMC.Med.Genomics.2011_PMID.21214954 | -0.0152 |
| miRNA_hsa-miR-425-3p | -0.0129 |
| Protein_PRAS40_pT246 | -0.0103 |
| Module_UNC_Duke_Module06_er_Median_Mike.PMID:20335537 | -0.0030 |
| Module_Pcorr_squamoid_PLOS.2012_PMID.22590557 | 0.0047 |
| Clinical_pathologic_N | 0.0069 |
| miRNA_hsa-miR-21-5p | 0.0079 |
| Module_UNC_MM_p53null.Basal_Median_Genome.Biol.2013_PMID.24220145 | 0.0104 |
| miRNA_hsa-miR-21-3p | 0.0119 |
| Protein_Caveolin-1 | 0.0138 |
| miRNA_hsa-miR-92b-3p | 0.0239 |
| Protein_TIGAR | 0.0275 |
| miRNA_hsa-miR-223-3p | 0.0287 |
| miRNA_hsa-miR-130a-3p | 0.0579 |
| miRNA_hsa-miR-222-3p | 0.0594 |
| Protein_IGFBP2 | 0.0643 |
| miRNA_hsa-let-7a-3p | 0.0700 |
| Clinical_age | 0.1147 |
| Module_UNC_Scorr_Basal_Correlation_JCO.2009_PMID.19204204 | 0.1352 |
| Clinical_pathologic_T | 0.2535 |

NOTE: See NOTE to Table 1.

**Table 3:**
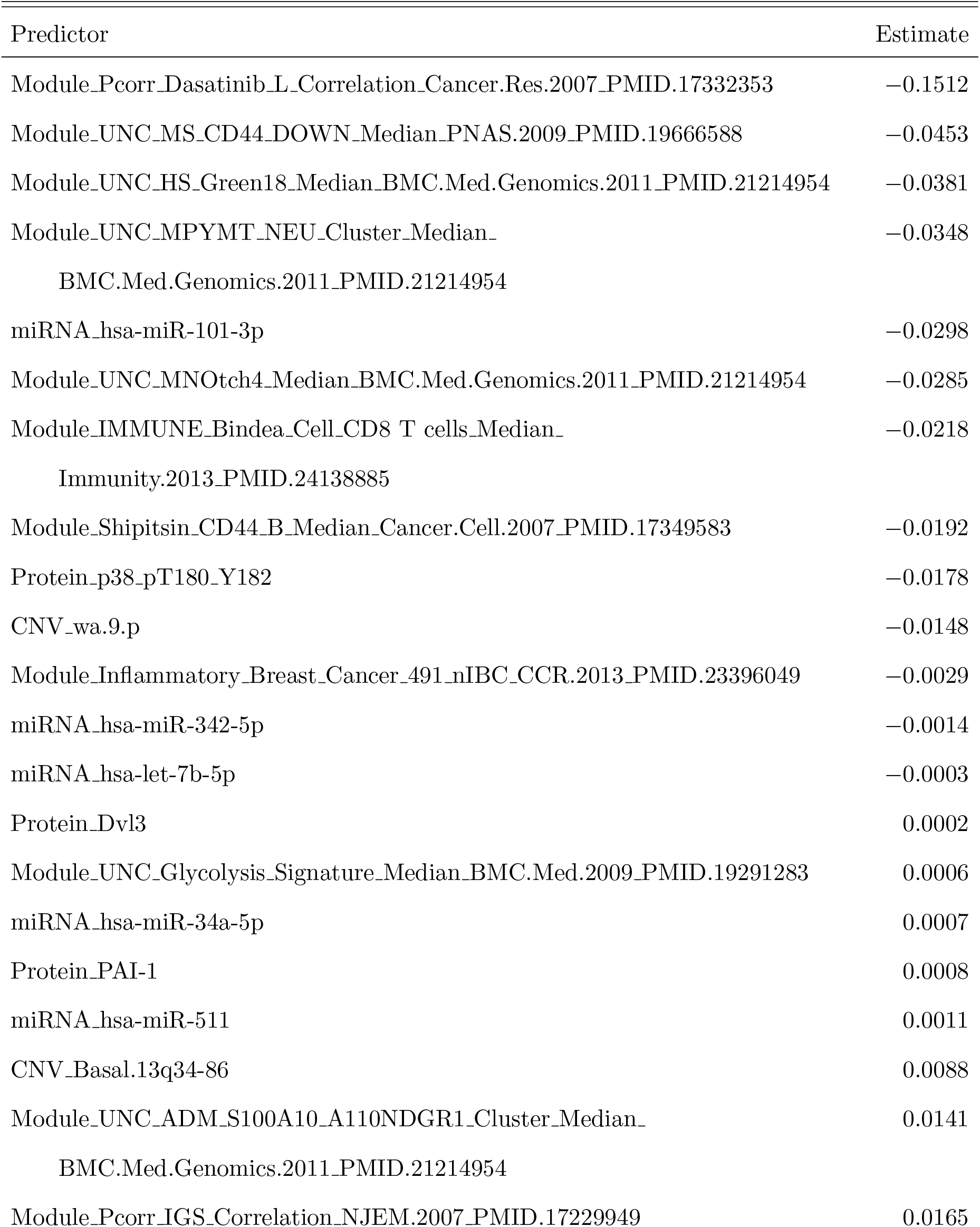
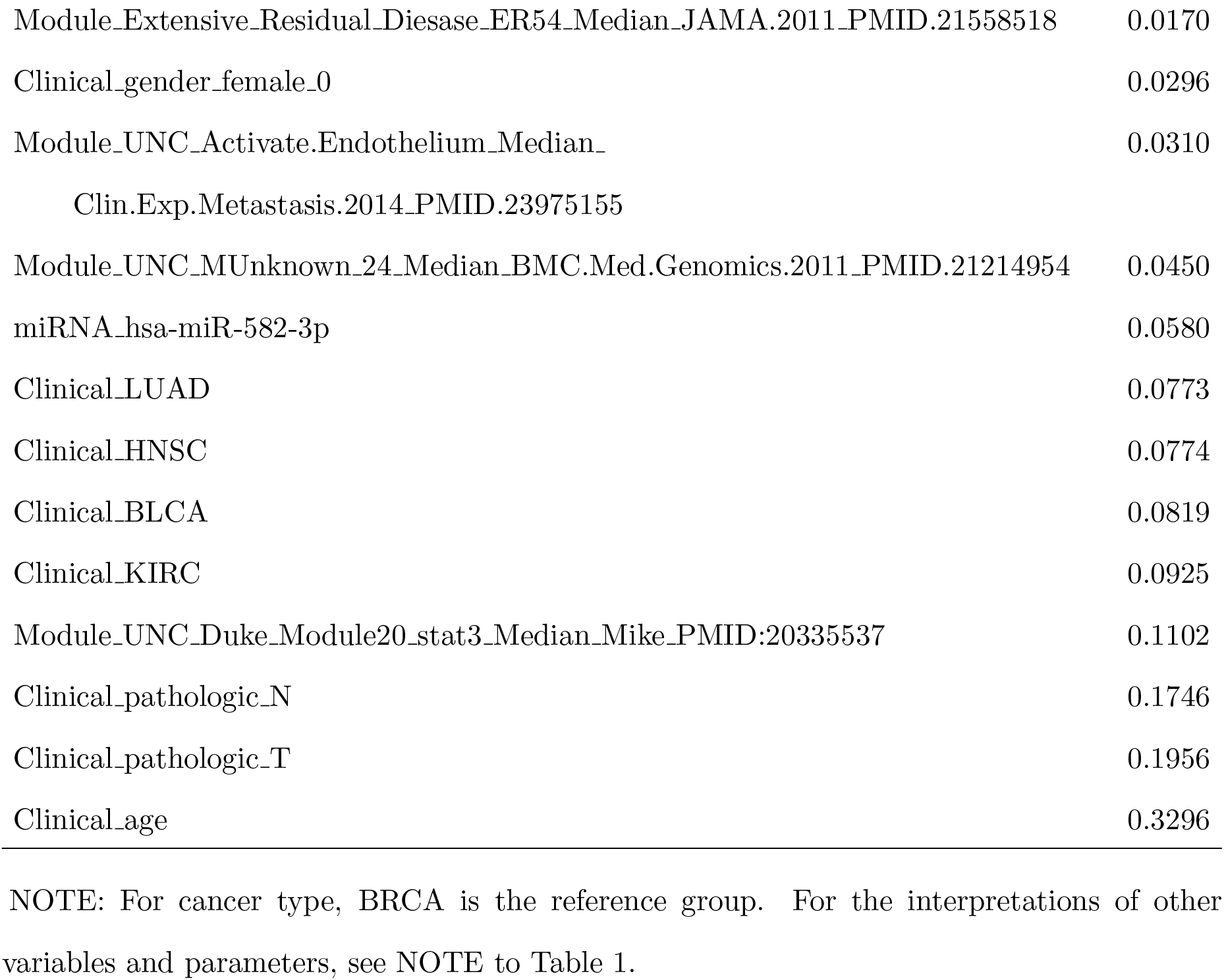
Analysis results from I-Boost-Permutation for the TCGA pan-cancer data set.

Age and pathological nodal status are negatively associated with survival time in the LUAD, KIRC, and pan-cancer data sets. Age has been reported to be prognostic for many cancer types [27, 28, 29]. In the analysis of the pan-cancer data set, cancer types were selected, which is logical, since the survival time is known to depend on cancer types [25]. Thus, the tissue of origin remains an important prognostic factor. Among the gene modules, Glycolysis_signature and MUnknown_24 are negatively associated with survival time in the LUAD and pan-cancer data sets; these two modules are correlated with Hypoxia signatures among a set of 1,198 TCGA breast cancer patients. Likewise, Pcorr_IGS_Correlation and Activate.Endothelium, which are negatively associated with survival time in the pan-cancer data set, are correlated with proliferation signatures; the latter are known to be negatively associated with survival time.

In contrast, signatures of CD8 T cells, non-inflammatory breast cancer (nIBC and MM_Red2), and luminal features (Mature_LuminalUp, GP7_estrogen signaling, HS_Green1, HS_Green8, LUMINAL_Cluster, Duke_Module06_er, Pcorr_Dasatinib_L_ Correlation, and HS_Green18) are positively associated with survival time in the KIRC or pan-cancer data sets. The NEU_cluster module is positively associated with survival time in the LUAD data set, which is biologically significant because this module represents epithelial luminal cell differentiation and thus tracks more differentiated and lower grade lung cancers. These selected features, together with their biological implications, demonstrate the robustness of the I-Boost methodology.

## Conclusions

In this paper, we present a novel method, termed I-Boost, for variable selection and outcome prediction that is especially powerful when one wishes to simultaneously consider multiple genomics and/or proteomics data types. We used simulation studies and real data to demonstrate that in the presence of multiple data types with diverse signal strength, I-Boost produces better outcome prediction than LASSO and elastic net. We proposed two versions of I-Boost, namely I-Boost-CV and I-Boost-Permutation. I-Boost-CV yields more accurate prediction than I-Boost-Permutation, but it generally selects many more variables and is computationally more intensive. By contrast, I-Boost-Permutation is computationally efficient and selects much fewer variables, which may be preferable for follow-up experiments.

Consistent with the current literature, we found that clinical variables are strong predictors of survival time. With I-Boost, we were able to build upon the clinical variables and extract additional useful information from genomic variables in order to improve the prediction; the improvement that we obtained with I-Boost was considerably larger than that obtained by either LASSO or elastic net. We also compared the use of individual gene expression data versus gene modules and found that the use of gene modules leads to improvement in prediction accuracy and more interpretable results. When we considered the selected I-Boost models, clinical variables (e.g., age, tumor size, and pathological nodal status) were strong predictors of survival. The I-Boost methods also selected several gene modules that were previously identified as prognostic of outcomes, whether positive or negative.

Our study has limitations. The main limitation is that the LUAD and KIRC data sets pertain to a relatively small number of patients, with an even smaller number of observed events. This limitation motivated us to combine eight solid epithelial tumor types to form a large pan-cancer data set. The analyses on the pan-cancer data might not properly account for heterogeneity across different cancer types. Another limitation of our study is that the quality of the clinical data varies across different cancer types; for example, the follow-up time for some cancer types was quite short.

In summary, we demonstrated that the performance of I-Boost is superior to that of elastic net and LASSO and that the performance of gene modules is superior to that of the totality of individual genes. The I-Boost methodology is applicable to any disease states where multiple types of genomics and/or proteomics data are available and thus has potential applications beyond cancer studies.

## Methods

### Data description

TCGA provides a large open-access database that includes clinical and genomics data for patients with 33 cancer types or subtypes. Herein, we focused on eight cancer types or subtypes, namely, LUAD, KIRC, colon adenocarcinoma (COAD), rectal adenocarcinoma (READ), lung squamous cell carcinoma (LUSC), bladder urothelial carcinoma (BLCA), breast invasive carcinoma (BRCA), and head and neck squamous cell carcinoma (HNSC). For clinical variables, somatic mutation, copy number variation, mRNA expression, and miRNA expression, data on 2,272 patients were obtained from the December 22, 2012 Pan-Cancer-12 data freeze from the Sage Bionetworks repository Synapse (http://www.synapse.org); the data were previously processed and described by Hoadley et al. [25]. Protein expression data were downloaded from Broad GDAC Firehose (http://gdac.broadinstitute.org/) on June 26, 2017 for a subset of 1,779 patients included in the data set of Hoadley et al. [25].

Clinical variables included gender, age, pathological stages T and N, and cancer type. In all analyses, COAD and READ were considered as one cancer type. For mRNA expression data, we used RNA-seq by Expectation-Maximization (RSEM) [30] to quantify the transcript abundances measured by RNA sequencing and used the log2-transformed up-quantile-normalized RSEM values of 12,434 genes. The RNA sequencing was performed at the University of North Carolina at Chapel Hill [31, 32, 33]. Gene level expression data are also available on the TCGA Data Portal (http://tcga-data.nci.nih.gov/tcga/). For mutation data, we used the single nucleotide variant calls, which were de-duplicated and re-annotated using the Ensembl version 69 transcript database. A total of 130 genes with non-synonymous mutations in more than 10% of the whole sample were included for the analyses. The combined mutation annotation format file is available from the Synapse resource. For miRNA expression data, we used the read count data for 305 normal ized expressions, which were compiled into an abundance matrix for 5p and 3p mature miRBase strands [31]. For reverse-phase protein arrays, we used the level-3 normalized data for 136 proteins or phospho-proteins. For copy number data, SNP6.0 array-based gene-level somatic copy number alteration data were generated from the GISTIC analysis [34]. The input data matrix is available in Synapse at syn1710678. We used the copy number values for 216 cancer-specific segments, which are frequently altered in cancer of various types including breast cancer, and segments for all chromosome arms (a total of 41 segments) [35, 36].

We defined gene modules as sets of co-expressed genes that are considered to be functional units in breast cancer. We built a collection of 497 gene modules. The modules were constructed on the basis of 73 publications or results from the Gene Set Enrichment Analysis [37]. A partial list of the modules appears in Fan et al. [12]. Among the modules, 461 are median expression values for homogeneously expressed genes, 33 are correlations of expression values with predetermined gene centroids, and 3 are built from previously published gene expression prognostic models.

After removing patients with missing data, the total sample size was 1,420, including 202 LUAD patients and 195 KIRC patients. All survival times were censored at five years if the patients were still in the study at that time point. For the pan-cancer data set, the median follow-up time was 16.8 months, and the censoring rate was 77.6%. For the subset of LUAD patients, the median follow-up time was 13.9 months, and the censoring rate was 71.3%. For the subset of KIRC patients, the median follow-up time was 28.9 months, and the censoring rate was 63.6%.

### LASSO and elastic net

We implemented LASSO and elastic net using the R-package “glmnet” [38] and used five-fold cross validation to select the tuning parameters. For elastic net, cross validation was performed over a two-dimensional grid of (*α*, λ), while for LASSO, *α* was set to be 1.

For elastic net, the grid for *α* was chosen to be (0.05, 0.1,0.2,…, 1.0), and a grid for λ was chosen separately for each *α* using the default settings of glmnet. (A minimum value of 0.05 was considered for *α*, because *α* too close to 0 may result in too many variables being selected; in particular, no sparsity is imposed if *α* = 0.) To make the selection procedure more stable, we repeated the split and evaluation procedure five times, and the cross-validation errors were averaged over the five repetitions.

### I-Boost

The I-Boost algorithm is given as follows:

1. Set *f*_0,*i*_ = 0 for *i* = 1,…, *n*, and let ***f***_0_ = (*f*_0,1_,…, *f*_0,*n*_)^T^.
2. Consider *m* = 1, 2,…:

a. For a given *k_m_* ∈ {1,…, *K*}, calculate

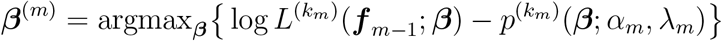

using the coordinate-descent algorithm [38], where

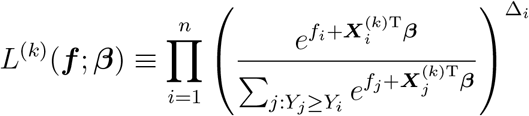

is the partial likelihood with offset term ***f*** and covariates ***X***^(*k*)^, *α_m_* and *λ_m_* are tuning parameters, ***f*** = (*f*_1_,…, *f_n_*)^T^, and 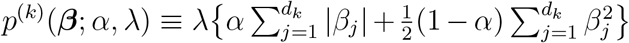 is the elastic net penalty. The selection of *k_m_*, *α_m_*, and *λ_m_* is described below.
b. Set 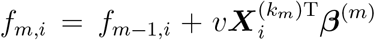 for *i* = 1,…, *n* with *v* = 0.1 and ***f***_*m*_ = (*f*_*m*,1_,…, *f_m,n_*)^T^.

At the *m*th iteration, only the regression parameters corresponding to the *k_m_*th data type are updated. We refer to the *d*-vector with value *β*^(*m*)^ at the positions corresponding to the *k_m_*th data type and zero elsewhere as the current estimate at the *m*th iteration. The current estimate at each iteration contributes to the final parameter estimate additively, and the final parameter estimate is simply the sum of the current estimates obtained from all steps multiplied by *v*.

I-Boost-CV and I-Boost-Permutation use cross validation and permutation, respectively, to choose (*k_m_*, *α_m_*, *λ_m_*) at step 2(a). For I-Boost-CV, we adopt five-fold cross validation separately at each iteration over a three-dimensional grid on 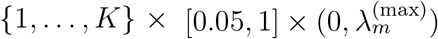 for (*k_m_*, *α_m_*, *λ_m_*), where 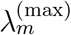 is a value large enough to shrink the current estimate to zero.

For I-Boost-Permutation, we first perform LASSO separately for each data type ***X***^(*k*)^ (*k* = 1,…, *K*) with tuning parameter 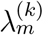, where 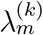 is selected using the permutation method proposed by Sabourin et al. [39]; the permutation method is only applicable to LASSO. The procedure is motivated by the principle that in a null model, i.e., in the absence of any relevant predictors, the tuning parameters should be chosen such that no variable is selected. The permutation selection procedure first generates hypothetical null models by randomly permuting (*Y_i_*, Δ_*i*_, *f*_*m*–1,*i*_) *B* times at each iteration, so that in each permuted data set the association between the predictors and the outcome (and the offset term) is removed. The procedure then finds the smallest *λ* such that no variable is selected for each permuted data set and selects the median of the *B* values of *λ*. For the *k*th data type (*k* = 1,…, *K*), let 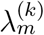 be the selected tuning parameter and 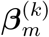 be the corresponding LASSO estimate. We select *k_m_* based on the partial-likelihood value at 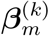, i.e., 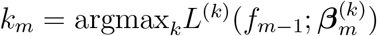, and set *α_m_* = 1 and 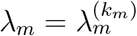.

Empirical studies suggested that a small value of the step-length factor *v* often improves and almost never worsens the performance of boosting [40]. Therefore, it is recommended that *v* is chosen to be as small as possible while the algorithm remains computationally feasible. In the settings we have considered, the performance of I-Boost is not sensitive to *v* within the range of *v* ∈ (0.05, 0.5). Therefore, we set *v* to a moderately small value of 0.1.

Conventional boosting methods require a stopping criterion to avoid over-fitting. In our experience, however, because the tuning parameters are selected separately at each iteration for I-Boost, they eventually lead to shrinkage of all (current) parameter estimates. Therefore, we do not adopt a separate procedure to determine the stopping time of the iteration. We terminate the iteration when ***f***_*m*_ remains constant for five consecutive iterations.

### Simulation studies

In the simulation studies, we considered all data types except individual gene expression data. For each simulation data set, we generated the predictors by sampling without replacement whole vectors of predictors from the TCGA pan-cancer data set. We generated the survival time from a proportional hazards model with the baseline hazard function *h*_0_(*t*) = *t* and generated the censoring time from an exponential distribution with a mean chosen to result in censoring proportion of about 50%. We set the sample size *n* to 500 in all settings.

The regression parameters were chosen to produce a different proportion of signals across data types, where the signal of data type *k* is defined to be 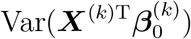, and the predictors were standardized. The variables with non-zero regression parameters, hereafter referred to as signal variables, were chosen to be weakly correlated. We considered three settings, with the distributions of signals and number of signal variables shown at the bottom of Figure 1. In all settings, the signals of all data types sum to 1.2, and the regression parameters of signal variables of the same data type are equal; based on simulation studies not presented, the relative performance of different methods is very similar under different values of total signal. In Setting 1, the clinical variables contain much stronger signals than the other data types. Mutation and copy number variation data do not contain any signal. In Setting 2, all signals are concentrated on the clinical variables and gene modules, and the two data types equally share the signals. In Setting 3, the clinical variables contain the most signals, and the remaining signals are evenly distributed across the other data types.

Because we considered a total of six data types, I-Boost-CV is computationally demanding. To lessen the computational burden, we set *v* = 0.2 instead of the value 0.1 used in real data analysis.

### Assessment of Prediction

To assess an analysis method, we split the data into 30 training and testing sets with a 3:2 ratio of sample sizes. We used the R-package “sampling” [41] to perform the data split, such that the distributions of the clinical variables in the training and testing sets are approximately equal. We performed the analysis on the training sets, and the results were assessed on the corresponding testing sets using the C-index. For each split of the data, we repeated this estimation-validation procedure on different combinations of data types as predictors. We only consider combinations of data types that include clinical variables, because clinical variables are almost always considered in practice, and one of the main objectives of this paper is to evaluate the prognostic value of the combination of genomics and clinical data. The analyses were conducted on the 30 splits of the data and on the 48 combinations of data types for the LUAD, KIRC, and pan-cancer data sets.

To quantify the prediction accuracy, we used the C-index. Let *T_i_* be the survival time and ***X***_*i*_ be a vector of predictors for the *i*th subject, and let *β* be a vector of regression parameters. The risk score is defined as 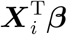. If *T_i_* and 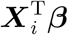 are continuous, then the C-index is defined as 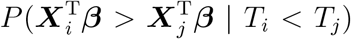. The C-index is the probability that for a random pair of subjects in which the first subject has a shorter survival time, the risk score for the first subject is higher. Thus, C-index measures how well the risk score aligns with the actual survival time. For each pair of training and testing sets, we set *β* to be the parameter estimate obtained from the training set and estimated the C-index for the testing set using the method of Pencina & D’Agostino [26]. If no variable was selected, then a C-index value of 0.5 was assigned.

## Declarations

### Authors’ contributions

DYL and CMP coordinated the overall studies. KYW performed the statistical analyses. KYW, ABN, DZ, and DYL developed the statistical methodologies. CF and JSP coordinated processing of the data. KYW, MT, DYL, and CMP wrote the paper, which all authors reviewed.

### Funding

This work was supported with funds from the NCI Breast SPORE program (P50-CA58223-09A1), the Breast Cancer Research Foundation, the Susan G. Komen, The V Foundation for Cancer Research, and by National Institutes of Health grants R01CA148761, R01GM047845, R01HG009974, and P01CA142538.

